# Analysis of structure-activity and structure-mechanism relationships among thyroid stimulating hormone receptor binding chemicals by leveraging ToxCast library

**DOI:** 10.1101/2023.06.14.544937

**Authors:** Ajaya Kumar Sahoo, Shanmuga Priya Baskaran, Nikhil Chivukula, Kishan Kumar, Areejit Samal

## Abstract

Thyroid stimulating hormone receptor (TSHR) is an integral part of the hypothalamic-pituitary-thyroid axis. Notably, dysregulation in TSHR activation in humans can lead to adverse effects such as Grave’s disease, hypothyroidism and Hashimoto’s disease. Moreover, animal studies have shown that binding of endocrine disrupting chemicals (EDCs) with TSHR can lead to developmental toxicity. Several such chemicals have also been screened for their adverse physiological effects in human cell lines through various high-throughput assays under the ToxCast project. The vast resource of data generated through ToxCast has enabled the development of different toxicity predictors, but they can be limited in their predictive ability due to the heterogeneity in structure-activity relationships among chemicals. In an attempt to explore this heterogeneity, we systematically investigated structure-activity and structure-mechanism relationships among the TSHR binding chemicals from ToxCast. By employing structure-activity similarity (SAS) map, we identified 79 activity cliffs among 509 chemicals in the TSHR agonist dataset and 69 activity cliffs among 650 chemicals in the TSHR antagonist dataset. Further, by using the matched molecular pair (MMP) approach, we find that the resultant activity cliffs (MMP-cliffs) are a subset of activity cliffs identified via the SAS map approach. Moreover, by leveraging ToxCast mechanism of action (MOA) annotations for chemicals common to both TSHR agonist and antagonist datasets, we identified 3 chemical pairs as Strong MOA-cliffs and 19 chemical pairs as Weak MOA-cliffs. In conclusion, the insights from this systematic analysis of the structure-activity as well as the structure-mechanism relationships of TSHR binding chemicals are likely to inform ongoing efforts towards development of better predictive toxicity models for characterizing the chemical exposome.

## 1. Introduction

Thyroid stimulating hormone receptor (TSHR) plays an important role in the hypothalamic-pituitary-thyroid axis where it mediates the production of thyroid hormone upon activation by the physiologic agonist, thyroid stimulating hormone (TSH) (Bassett and Williams, 2008; Feldt-Rasmussen et al., 2021; Ortiga-Carvalho et al., 2016). The hypothalamic-pituitary-thyroid axis is crucial for development and metabolism, and is prone to disruptions by endocrine disrupting chemicals (EDCs) (Fekete and Lechan, 2014; Schmutzler Cornelia et al., 2007; Thambirajah et al., 2022) in the human exposome. In particular, animal studies have shown that EDCs binding to TSHR disrupt the thyroid system, ultimately leading to developmental toxicity (Chen et al., 2020; Lee et al., 2022; Teng et al., 2018). In human, the overproduction of thyroid hormone caused by the binding of M22 autoantibody with TSHR can lead to Grave’s disease (Sanders et al., 2003), and underproduction of thyroid hormone caused by the binding of K1-70 autoantibody can lead to hypothyroidism and Hashimoto’s disease (Evans et al., 2010). Consequently, screening of environmental chemicals in the human exposome that can bind to TSHR is important for their proper management.

The assessment of adverse effects of environmental chemicals on physiological targets is a laborious, time-consuming process and might involve animal testing. In this direction, the ToxCast program has screened nearly 10,000 chemicals for their adverse effects on various biological targets including TSHR, and characterized them based on their bioactivity and mechanisms of action (Richard et al., 2021, 2016). The ToxCast dataset has greatly enabled the development of several quantitative structure-activity relationship (QSAR) models that aim to predict toxicity of chemicals and aid in prioritization of chemicals for further testing (Dix et al., 2007; Jeong et al., 2022). In particular, the ToxCast library has been used to develop machine learning based QSAR models to predict chemicals that bind to TSHR (Garcia de Lomana et al., 2021; Kurosaki et al., 2020). However, the heterogeneity of the structure-activity landscape of chemicals that bind to TSHR has not been explored while developing such models, which could lead to uncertainties in associated predictions (Mathea et al., 2016).

The heterogeneity in the structure-activity landscape of chemicals arises due to the presence of activity cliffs (Cruz-Monteagudo et al., 2014). Activity cliffs are formed by chemical pairs that have similar structures but significantly differ in their activity values (Stumpfe et al., 2019). The identification of activity cliffs in a chemical dataset is necessary as it limits the predictive power of QSAR models (Maggiora, 2006). Many methods have been developed for the analysis of the structure-activity landscape of chemicals and identification of activity cliffs (Guha and Van Drie, 2008; Hu and Bajorath, 2012; Medina-Franco et al., 2009; Peltason and Bajorath, 2007; Wawer et al., 2008). Medina-Franco and colleagues have extensively used the chemical fingerprint-based structure-activity similarity (SAS) map to identify activity cliffs in diverse chemical datasets (Méndez-Lucio et al., 2012; Naveja et al., 2018; Naveja and Medina-Franco, 2015). In an earlier contribution, some of us had extended this approach to identify and characterize activity cliffs in androgen receptor binding chemicals (Vivek-Ananth et al., 2023). Independently, Bajorath and colleagues have developed a substructure-based matched molecular pair (MMP) approach to identify activity cliffs (Hu et al., 2012). This approach had been extended by Hao *et al*. (Hao et al., 2016) to identify the differences in the mechanisms of action of chemical pairs with similar structures, and moreover, introduced the concept of mechanism of action cliffs (MOA-cliffs). Like activity cliffs, the presence of MOA-cliffs highlights the heterogeneity in the structure-mechanism relationships among chemicals. Importantly, an exploration of the heterogeneity in the structure-activity landscape in conjunction with the structure-mechanism relationships has not been conducted on the ToxCast chemical library to date, in particular, for the chemicals that can bind to TSHR.

In this study, we performed a systematic investigation of the structure-activity landscape and structure-mechanism relationships in datasets of TSHR agonist and TSHR antagonist compiled from ToxCast chemical library. We employed both SAS map and MMP based approaches to identify the activity cliffs in the structure-activity landscape of these chemical datasets. We classified the identified activity cliffs into different categories using the information on their chemical structures. Further, we leveraged the mechanism of action (MOA) annotations for chemicals common to both TSHR agonist and TSHR antagonist datasets to identify MOA-cliffs. To the best of our knowledge, we present the first systematic study leveraging ToxCast chemical library and employing multiple cheminformatics approaches for the identification and characterization of activity cliffs along with MOA-cliffs among chemicals that can bind to TSHR.

## 2. Materials and Methods

### 2.1. Chemical dataset comprising of agonists and antagonists of the thyroid stimulating hormone receptor

The objective of this investigation is the analysis of the structure-activity landscape of the agonists and antagonists of the thyroid stimulating hormone receptor (TSHR) (Figure 1). For this investigation, we retrieved the chemicals, their corresponding activity values, and endpoints from Tox21 assays (assay source identifier 7) within ToxCast version 3.5 (EPA, 2023) using level 5 and 6 processing. First, we used an in-house R script to filter the Tox21 multi-concentration summary file in order to identify chemicals based on their endpoint being either TSHR agonist (assay endpoint identifier 2040) or TSHR antagonist (assay endpoint identifier 2043) screened in HEK293T cell line. TSHR agonist is a chemical that binds to TSHR and fully activates it, whereas TSHR antagonist is a chemical that binds to TSHR but does not activate it and can additionally block the activation by any other agonist. Next, we filtered chemicals annotated as representative samples (i.e., gsid_rep is 1) and with reported activity value (i.e., modl_ga value is present) (Figure 1a). Subsequently, for these shortlisted chemicals, we accessed the two-dimensional (2D) structures provided by ToxCast version 3.5, or PubChem (https://pubchem.ncbi.nlm.nih.gov/), if the 2D structures were not provided by ToxCast. Thereafter, we used MayaChemTools (Sud, 2016) to remove salts, mixtures, invalid structures and duplicated chemicals (Figure 1a). We also removed linear chemicals using the scaffold definition employed in our previous work (Vivek-Ananth et al., 2023). Finally, we curated a TSHR agonist dataset of 509 chemicals (SI Table S1) and a TSHR antagonist dataset of 650 chemicals (SI Table S2). For each chemical in the two datasets, we additionally compiled the Chemical Abstracts Service (CAS) registry number or PubChem compound identifiers, reported biological activity (i.e., either active: hit_c is 1; or inactive: hit_c is 0), and the chemical concentration that generates the half maximal response (modl_ga, i.e., logarithm of AC_50_ values in micromolar concentration).

**Figure 1:**
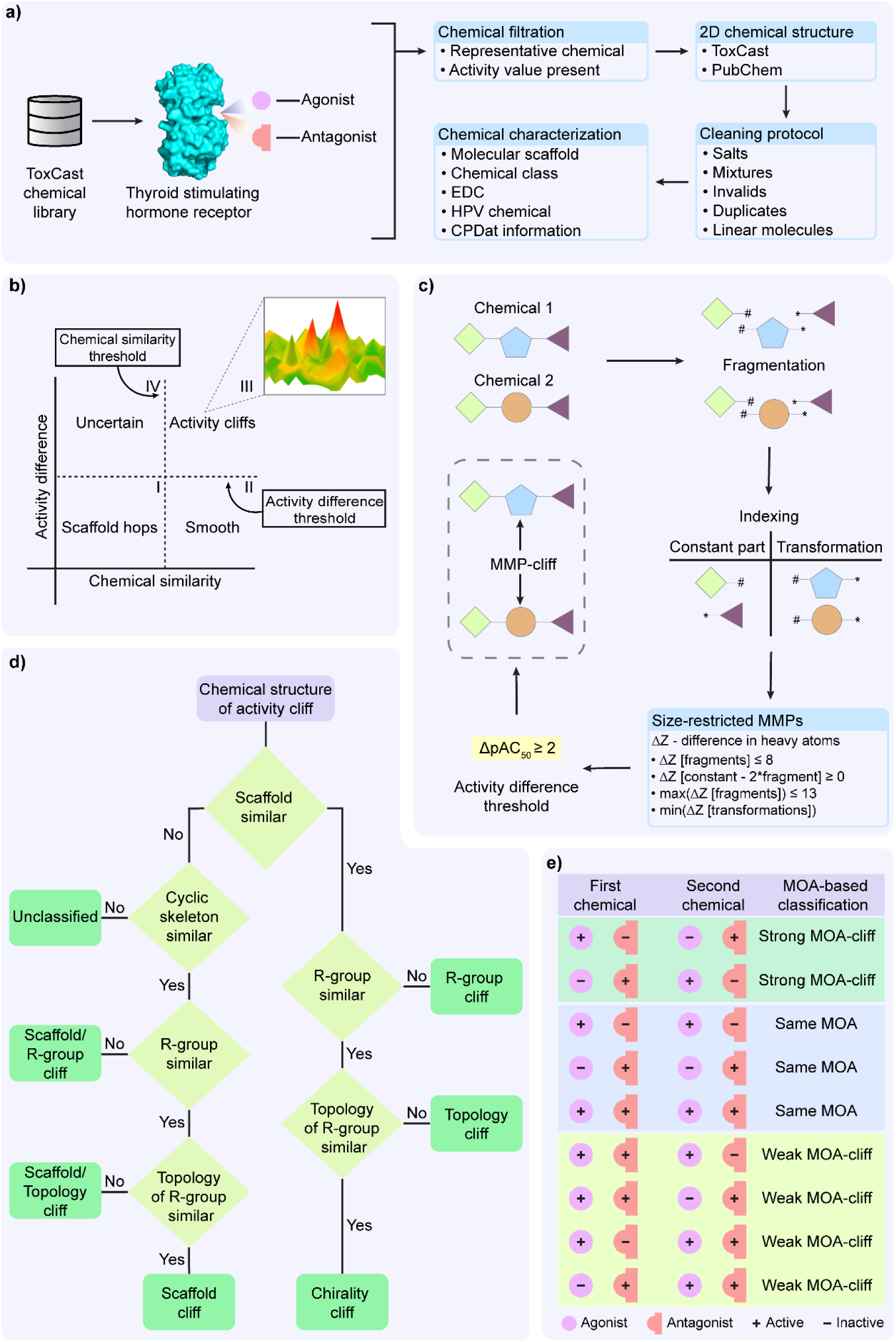
Summary of structure-activity landscape analysis and activity cliff identification in a chemical dataset curated from ToxCast library. **(a)** Curation and chemical characterization of thyroid stimulating hormone receptor (TSHR) agonist and antagonist datasets. **(b)** Structure-activity similarity (SAS) map based approach to identify the activity cliffs in a chemical dataset. **(c)** Steps involved in generation of a matched molecular pair (MMP) and associated MMP-cliff. **(d)** Classification of activity cliff pairs based on respective structural information. **(e)** Mechanism of action (MOA) based classification of the chemical pairs (common to both TSHR agonist and antagonist datasets and having Tanimoto coefficient based similarity of > 0.35) into three different categories.

### 2.2. Chemical characterization and annotation

We characterized chemicals in both TSHR agonist and TSHR antagonist datasets using molecular scaffolds and chemical classifications (Figure 1a). Following our previous work (Vivek-Ananth et al., 2023), we used the Bemis-Murcko definition (Bemis and Murcko, 1996) to compute the molecular scaffolds from chemical structures. Next, we relied on ClassyFire (Djoumbou Feunang et al., 2016) to provide the corresponding chemical classifications. Further, we used DEDuCT (Karthikeyan et al., 2021, 2019) to identify endocrine disrupting chemicals (EDCs), and Organisation for Economic Co-operation and Development High Production Volume (OECD HPV) (https://www.oecd.org/chemicalsafety/risk-assessment/33883530.pdf) or United States High Production Volume (USHPV) (https://comptox.epa.gov/dashboard/chemical-

<u>lists/EPAHPV</u>) databases to identify high production volume chemicals in our datasets. Additionally, we leveraged the CAS identifiers of the chemicals in TSHR agonist and TSHR antagonist datasets (which are also compiled in Distributed Structure-Searchable Toxicity (DSSTox) database) to retrieve annotations such as functional uses and occupational health hazard reports from Chemical and Products Database (CPDat) (Figure 1a) (Dionisio et al., 2018).

### 2.3. Computation of activity difference

The activity difference for a pair of chemicals is considered as the difference between their corresponding pAC_50_ values, where pAC_50_ is the negative logarithm of AC_50_ value in molar concentration (Hao et al., 2016; Méndez-Lucio et al., 2012; Pérez-Villanueva et al., 2011). The activity values of the chemicals in the compiled TSHR agonist and TSHR antagonist datasets are reported as the logarithm of AC_50_ values in micromolar concentrations (modl_ga). Therefore, we converted the modl_ga value to pAC_50_ value using the following formulae:

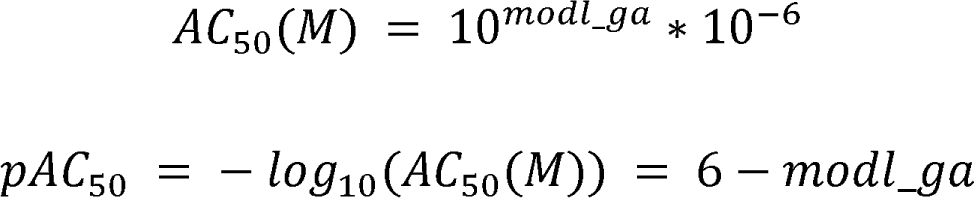

Thereafter, we calculated the activity difference between two chemicals *i* and *j* using the following formula:

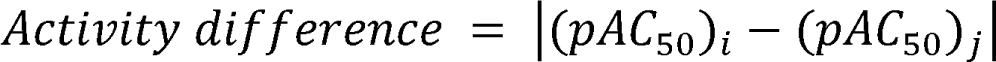

wherein the (pAC_SO_)_i_ and (pAC_SO_)_j_ are the pAC_SO_ values of chemicals *i* and *j* respectively.

### 2.4. Identification of activity cliffs using structure-activity similarity (SAS) map

We independently analyzed the activity landscape of the chemicals in TSHR agonist and TSHR antagonist datasets using structure-activity similarity (SAS) map (Figure 1b) (Méndez-Lucio et al., 2012; Naveja et al., 2018; Naveja and Medina-Franco, 2015; Vivek-Ananth et al., 2023). SAS map is a 2D representation where the structural similarity between the chemicals is plotted along the x-axis and the activity difference between the chemicals is plotted along the y-axis (Figure 1b). We computed structural similarity between chemical pairs based on Tanimoto coefficient between the corresponding Extended-Connectivity Fingerprints with diameter 4 (ECFP4) of the chemicals. As there is no strict rule to choose a threshold for high structural similarity (Medina-Franco, 2012), we considered a similarity threshold of 0.35 which was close to three standard deviations from median of the computed Tanimoto coefficient for chemical pairs in both TSHR agonist and TSHR antagonist datasets. We considered an activity difference threshold of 100 fold change which is equivalent to 2 logarithmic units (Naveja et al., 2018; Vivek-Ananth et al., 2023). Finally, we designated the highly similar chemical pairs (Tanimoto coefficient > 0.35) with high activity difference (≥ 2) as the activity cliffs in both TSHR agonist and TSHR antagonist datasets (Region III in Figure 1b). Additionally, we considered chemicals which form at least 5 activity cliff pairs as activity cliff generators (ACGs) (Naveja et al., 2018; Vivek-Ananth et al., 2023).

### 2.5. Identification of activity cliffs based on Matched Molecular Pairs (MMPs)

In addition to SAS map based activity landscape analysis, we employed the matched molecular pairs (MMP) based approach to identify the activity cliffs (MMP-cliffs) (Hu et al., 2012) independently in TSHR agonist and TSHR antagonist datasets (Figure 1c). We used mmpdb platform (Dalke et al., 2018) to generate MMPs for chemicals in both datasets. First, the mmpdb fragment module was used to fragment the chemical structure with ‘none’ value for both maximum number of non-hydrogen atoms and maximum number of rotatable bonds arguments. Next, the mmpdb index module was used to generate an exhaustive list of MMPs with ‘none’ value for maximum number of non-hydrogen atoms in the variable fragment argument. This gave us an exhaustive list of MMPs with detailed information on the constant part and transformations containing the exchanged fragments between chemical pairs. Further, to generate size-restricted MMPs, we implemented the following four criteria (Figure 1c) (Hu et al., 2012):

i. The difference in number of heavy atoms of the exchanged fragments in transformation should not exceed 8.
ii. The constant part should be at least twice the size of each fragment in the transformation.
iii. The number of heavy atoms of each fragment in the transformation should not exceed 13.
iv. For a chemical pair with multiple MMPs, the transformation with the least difference in the number of heavy atoms between the exchanged fragments is considered.

Finally, we identified MMP-cliffs among the size-restricted MMPs by selecting those pairs with an activity difference ≥ 2 in logarithmic units (i.e., 100 fold change) (Figure 1c).

### 2.6. Activity cliff classification

In this study, we followed the activity cliff classification described in Vivek-Ananth *et al*. (Vivek-Ananth et al., 2023), to classify the activity cliffs by considering their molecular scaffolds, R-groups, R-group topology and chirality of chemical structures. Here, we modified the workflow in Vivek-Ananth *et al*. (Vivek-Ananth et al., 2023) to also check for topologically equivalent scaffolds (cyclic skeleton) when a pair of chemicals do not share the same scaffolds (Figure 1d) (Hu and Bajorath, 2012). We used the R-group decomposition module available in RDKit (RDKit, 2023) to decompose the chemical structure into its core structure (scaffold) and R-groups. Further, we used the modified workflow (Figure 1d) to manually classify the activity cliffs into the following 7 types:

i. Chirality cliff: chemical pairs having the same scaffold, R-groups and R-group topology
ii. Topology cliff: chemical pairs having different R-group topologies while their scaffolds and R-groups remain unchanged
iii. R-group cliff: chemical pairs having different R-groups while their scaffolds remain unchanged
iv. Scaffold cliff: chemical pairs having different scaffolds while their cyclic skeletons, R-groups and R-group topologies remain unchanged
v. Scaffold/Topology cliff: chemical pairs having different scaffolds and R-group topologies while their cyclic skeletons and R-groups remain unchanged
vi. Scaffold/R-group cliff: chemical pairs having different scaffolds and R-groups while their cyclic skeletons remain unchanged
vii. Unclassified: chemical pairs having different scaffolds and cyclic skeletons

### 2.7. Mechanism of action (MOA) based classification of chemical structures

In addition to activity cliffs in TSHR agonist and TSHR antagonist datasets, we were interested in identifying chemical pairs in which the chemicals have similar structures but differ in their mechanism of action (MOA). Such chemical pairs are designated as MOA-cliffs (Hao et al., 2016). We considered chemicals which were common to both TSHR agonist and TSHR antagonist datasets, and removed those chemicals which were reported as inactive MOA in both assays. We then computed the structural similarity of chemical pairs by using the Tanimoto coefficients between the ECFP4 fingerprints of the shortlisted chemicals. We chose 0.35 as the similarity threshold (which is the structural similarity threshold used in SAS map analysis) to filter similar chemical pairs. Based on their MOA annotations in TSHR agonist and TSHR antagonist datasets, we classified these chemical pairs into 3 types (Figure 1e):

i. Strong MOA-cliff: chemical pairs in which the chemicals have opposite MOA annotations
ii. Same MOA: chemical pairs in which both the chemicals have same MOA annotations
iii. Weak MOA-cliff: chemical pairs which could not be classified as either Strong MOA-cliff or Same MOA

## 3. Results

### 3.1. Exploration of the chemical space of TSHR agonist and antagonist datasets

From ToxCast library, we curated 509 chemicals in TSHR agonist (SI Table S1) and 650 chemicals in TSHR antagonist (SI Table S2) datasets, and thereafter, annotated the chemicals in the two datasets with information on their molecular scaffolds, chemical classifications, and their presence in public documentation or databases (Methods; Figure 1a). Notably, there were 89 chemicals common between TSHR agonist and TSHR antagonist datasets.

For the 509 chemicals in the TSHR agonist dataset, after computing the molecular scaffolds we observed that the benzene scaffold is highly represented (as it is found in 122 chemicals). Many of the chemicals in TSHR agonist dataset are also categorized under the chemical class of ‘Benzene and substituted derivatives’ (195 chemicals) (SI Table S1). Importantly, 79 chemicals in the TSHR agonist dataset are documented in DEDuCT (Karthikeyan et al., 2021, 2019) as endocrine disrupting chemicals (EDCs) with experimental evidence, of which 21 chemicals are documented as high production volume chemicals as per OECD HPV or USHPV databases (Methods; SI Table S1). Chemical and Products Database (CPDat) provided various functional use annotations for 102 chemicals, of which biocides, fragrance and antioxidants are the major reported functional categories (SI Table S1). Additionally, 4 chemicals namely, 3-Carene, Butylated hydroxytoluene, Hydroquinone and Triphenyl phosphate have been documented in various occupational health hazard reports (SI Table S1).

Similarly, for the 650 chemicals in the TSHR antagonist dataset, we observed that benzene scaffold is the most represented molecular scaffold (as it is found in 127 chemicals), while ‘Benzene and substituted derivatives’ is the most represented chemical class (254 chemicals) (SI Table S2). Notably, 65 chemicals in the TSHR antagonist dataset are documented as EDCs in DEDuCT, of which 13 are also documented as high production volume chemicals in OECD HPV or USHPV databases (SI Table S2). CPDat provided functional uses for 156 chemicals, of which biocides, fragrance and antioxidants are reported as the major functional categories (SI Table S2). Additionally, 4 antagonists namely, 2,2’,4,4’,5-Pentabromodiphenyl ether, 2,2’,4,4’-Tetrabromodiphenyl ether, Bibenzyl and Styrene are documented in various occupational health hazard reports (SI Table S2).

### 3.2. Activity landscape analysis of TSHR agonist dataset

The structure-activity similarity (SAS) map has been employed in the literature to identify activity cliffs by investigating the structure-activity relationship (Méndez-Lucio et al., 2012; Naveja et al., 2018; Naveja and Medina-Franco, 2015; Vivek-Ananth et al., 2023). Accordingly, we analyzed the activity landscape of the TSHR agonist dataset using the SAS map approach (Methods; Figure 2a). We observed that the majority of chemical pairs show similar activity while they are structurally diverse (see SAS map Region 1 in Figure 2a). Importantly, we identified 79 chemical pairs showing high activity difference while being structurally similar (see SAS map Region III in Figure 2a). We designated these 79 chemical pairs (formed by 60 unique chemicals) as activity cliffs (SI Table S3), of which 9 chemicals are additionally identified as activity cliff generators (ACGs) (Methods; SI Table S4). The chemicals forming activity cliffs are represented by 34 unique scaffolds with benzene and triphenyltin scaffolds being the highly represented scaffolds, and are categorized under 13 chemical classes with ‘Benzene and substituted derivatives’ class being the largest category. Moreover, triphenyltin scaffold is highly represented in chemicals forming ACGs.

**Figure 2:**
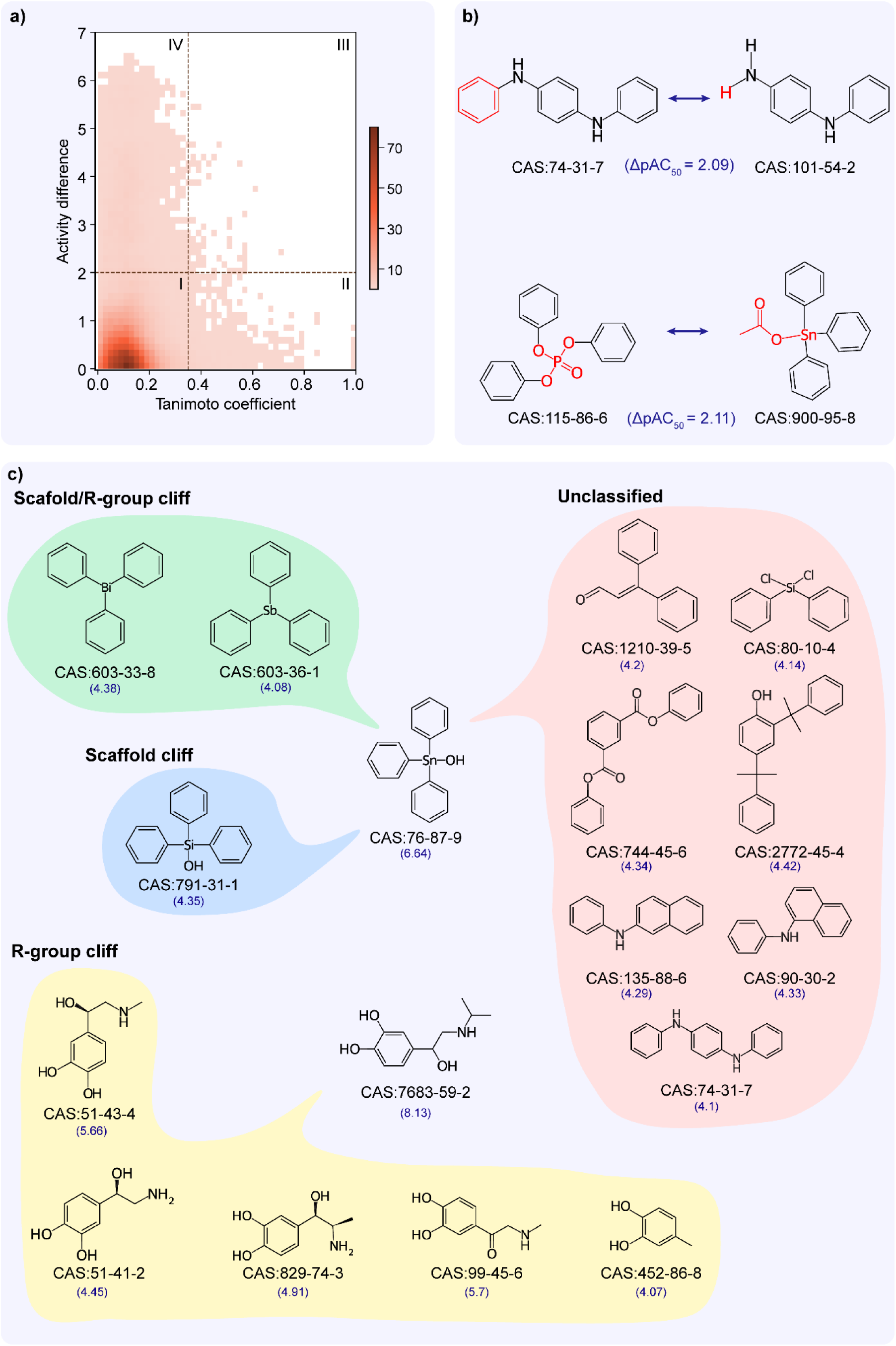
Activity landscape analysis of TSHR agonist dataset. **(a)** Structure-activity similarity (SAS) map for TSHR agonist dataset. SAS map is divided into 4 quadrants by considering a similarity threshold of 0.35 and activity difference threshold of 2. Further, the density of data points in different regions of the SAS map is shown using a color gradient. **(b)** MMP-cliffs formed by N,N’-Diphenyl-p-phenylenediamine (CAS identifier 74-31-7) with N-Phenyl-1,4-benzenediamine (CAS identifier 101-54-2) [LpAC_50_ = 2.09] and Triphenyl phosphate (CAS identifier 115-86-6) with Triphenyltin acetate (CAS identifier 900-95-8) [LpAC_50_ = 2.11]. The transformed fragments resulting in MMP-cliff are highlighted in red color. **(c)** Activity cliff classifications for the activity cliff generators, Triphenyltin hydroxide (CAS identifier 76-87-9; 10 activity cliff pairs) and Isoproterenol (CAS identifier 7683-59-2; 5 activity cliff pairs). The activity value (pAC_50_) is mentioned below for each chemical.

Matched Molecular Pair (MMP) based activity landscape analysis has been alternatively employed in the literature to identify the activity cliffs (Hao et al., 2016; Hu et al., 2012). We also used the MMP approach to analyze the activity landscape of the TSHR agonist dataset. We identified 523 MMPs formed by 170 chemicals in the TSHR agonist dataset (Methods; SI Table S5), of which 38 MMPs (formed by 19 unique chemical pairs) are identified as MMP-cliffs based on an activity difference threshold consideration similar to SAS map (Methods; SI Table S3). Notably, the MMP-cliffs identified by the MMP approach are a subset of the activity cliffs identified by the SAS map approach. Interestingly, the constant part containing three benzene rings identified in 14 of the 38 MMP-cliffs is similar to the highly represented triphenyltin scaffold among the chemicals forming activity cliffs identified through SAS map. Figure 2b shows chemical pairs of N,N’-Diphenyl-p-phenylenediamine (CAS identifier 74-31-7) and N-Phenyl-1,4-benzenediamine (CAS identifier 101-54-2), Triphenyl phosphate (CAS identifier 115-86-6) and Triphenyltin acetate (CAS identifier 900-95-8) that are identified as MMP-cliffs. N,N’-Diphenyl-p-phenylenediamine is an ACG which is documented as an EDC in DEDuCT and present in the OECD HPV or USHPV databases. Notably, Triphenyl phosphate and Triphenyltin acetate are documented as EDCs in DEDuCT and Triphenyl phosphate is also present in the OECD HPV or USHPV databases.

Subsequently, we classified the 79 activity cliffs and identified 11 as R-group cliffs, 1 as scaffold cliff, 11 as Scaffold/R-group cliffs and 56 as unclassified (Methods; SI Table S3). Figure 2c shows the different classifications of the activity cliffs formed by N,N’-Diphenyl-p-phenylenediamine and Isoproterenol (CAS identifier 7683-59-2). N,N’-Diphenyl-p-phenylenediamine forms 10 activity cliff pairs where 2 are Scaffold/R-group cliffs, 1 is Scaffold cliff and remaining are Unclassified. Similarly, Isoproterenol forms 5 activity cliff pairs where all are R-group cliffs.

### 3.3. Activity landscape analysis of TSHR antagonist dataset

Similar to the activity landscape analysis of the TSHR agonist dataset, we analyzed the TSHR antagonist dataset through both SAS map and MMP approaches. From the SAS map approach, while most chemical pairs show similar activity despite having diverse structures (see SAS map Region I in Figure 3a), 69 chemical pairs showed high activity difference while they are structurally similar (see SAS map Region III in Figure 3a). We designated these 69 chemical pairs as activity cliffs, and observed that they are formed by 75 unique chemicals (SI Table S6), of which 4 chemicals are ACGs (Methods; SI Table S7). The chemicals forming activity cliffs are represented by 39 unique scaffolds with benzene and biphenyl scaffolds being the highly represented scaffolds, and are categorized under 17 chemical classes with ‘Benzene and substituted derivatives’ class being the largest category. From the MMP approach, we identified 590 MMPs (formed by 195 chemicals), of which 3 are MMP-cliffs (Methods; SI Table S8).

**Figure 3:**
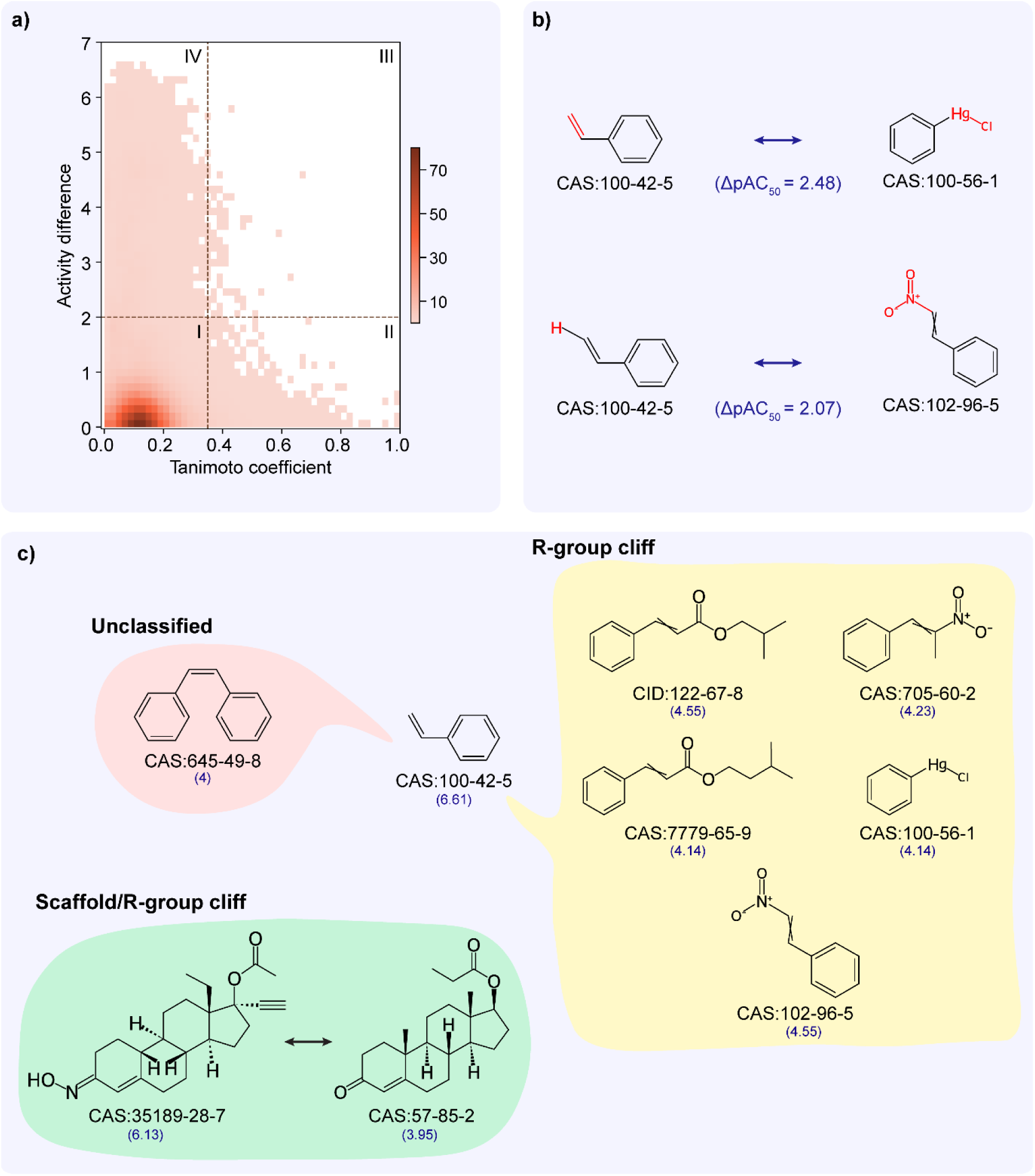
Activity landscape analysis of TSHR antagonist dataset. **(a)** Structure-activity similarity (SAS) map for TSHR antagonist dataset. SAS map is divided into 4 quadrants by considering a similarity threshold of 0.35 and activity difference threshold of 2. Further, the density of data points in different regions of the SAS map is shown using a color gradient. **(b)** MMP-cliffs formed by Styrene (CAS identifier 100-42-5) with Phenylmercuric chloride (CAS identifier 100-56-1) [LpAC_50_ = 2.48] and with beta-Nitrostyrene (CAS identifier 102-96-5) [LpAC_50_ = 2.07]. The transformed fragments resulting in MMP-cliff are highlighted in red color. **(c)** Activity cliff classifications for the activity cliff generator, Styrene (6 activity cliff pairs) and an activity cliff pair of Norgestimate (CAS identifier 35189-28-7) with Testosterone propionate (CAS identifier 57-85-2). The activity value (pAC_50_) is mentioned below for each chemical.

Notably all the MMP-cliffs are also activity cliffs identified through SAS map approach. Figure 3b shows chemical pairs of Styrene (CAS identifier 100-42-5) and Phenylmercuric chloride (CAS identifier 100-56-1), and Styrene and beta-Nitrostyrene (CAS identifier 102-96-5). Styrene is an ACG which is documented as an EDC in DEDuCT and present in the OECD HPV or USHPV databases.

Further, we classified the 69 activity cliffs and identified 18 as R-group cliffs, 1 as Scaffold/R-group cliff and 50 as Unclassified (Methods; SI Table S6). Figure 3c shows 6 activity cliffs formed by Styrene (5 R-group cliffs and 1 Unclassified) and 1 Scaffold/R-group cliff formed by Norgestimate (CAS identifier 35189-28-7) and Testosterone propionate (CAS identifier 57-85-2).

### 3.4. Mechanism of action (MOA) cliffs

Apart from the differences in activity, chemicals also show a difference in their identified mechanism of action (MOA). Hao *et al*. (Hao et al., 2016) have earlier explored the MMPs with different MOAs from androgen receptor agonist and antagonist datasets, and designated them as MOA-cliffs. We shortlisted 75 chemicals which have endpoints in both TSHR agonist and TSHR antagonist datasets and identified 38 chemical pairs which have high structural similarity (Methods; SI Table S9). We classified these 38 chemical pairs based on their MOA annotations and identified 3 as Strong MOA-cliffs, 16 as Same MOA and 19 as Weak MOA-cliffs (Methods; Figure 1e; SI Table S9). Notably, 1 Strong MOA-cliff and 8 Weak MOA-cliffs are also classified as activity cliffs identified through the SAS map approach. Figure 4 shows examples of different MOA based classifications of chemical pairs. 3,3’-Diaminobenzidine (CAS identifier 91-95-2) and 3,3’-Dimethylbenzidine (CAS identifier 119-93-7) form Strong MOA-cliff, Triphenyltin chloride (CAS identifier 639-58-7) and Triphenyltin hydroxide (CAS identifier 76-87-9) form Same MOA, and Endosulfan sulfate (CAS identifier 1031-07-8) and Endosulfan I (CAS identifier 959-98-8) form Weak MOA-cliff.

**Figure 4:**
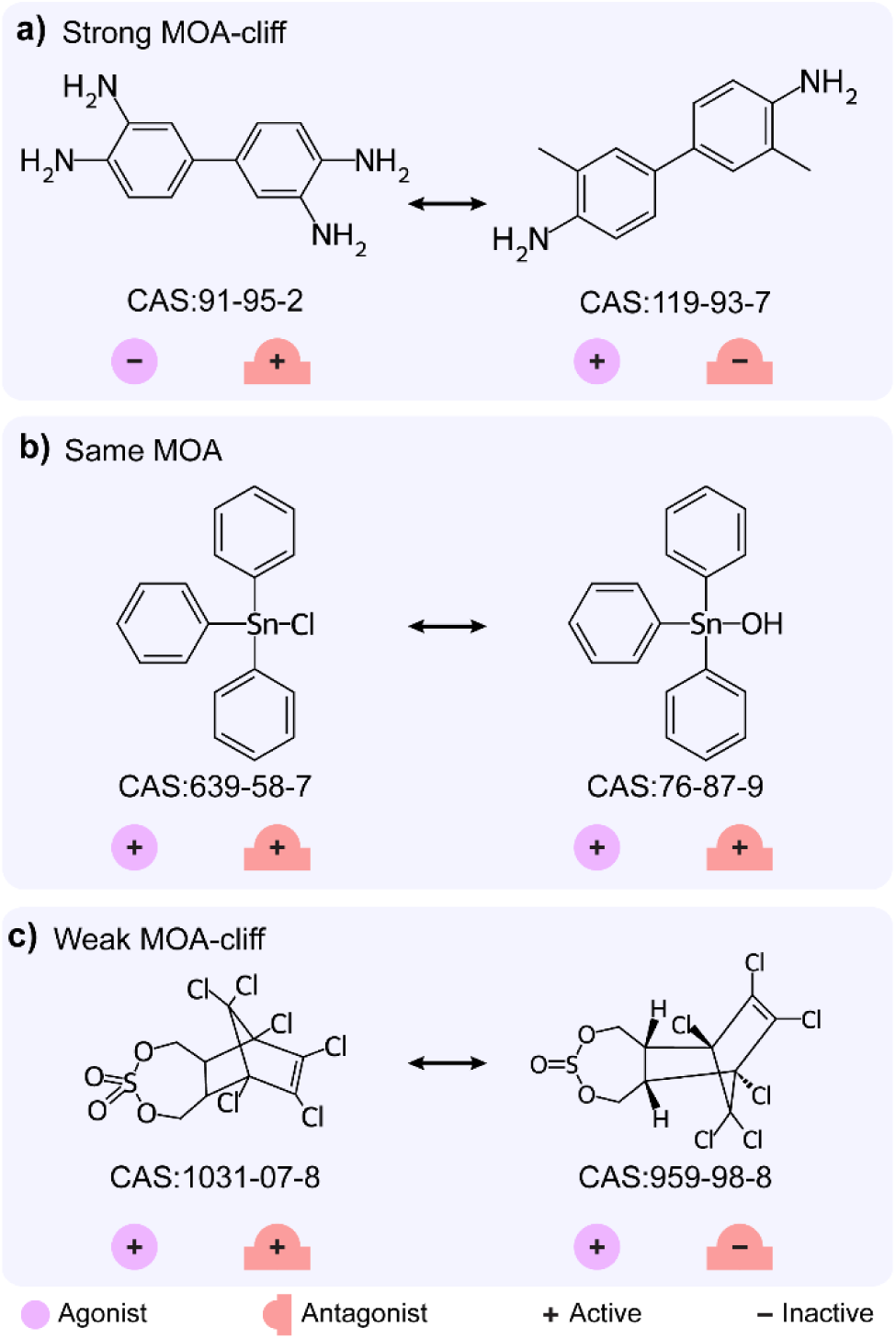
Examples for three different mechanism of action (MOA) based classifications of chemical pairs. **(a)** Strong MOA-cliff formed by 3,3’-Diaminobenzidine (CAS identifier 91-95-2) with 3,3’-Dimethylbenzidine (CAS identifier 119-93-7). **(b)** Same MOA formed by Triphenyltin chloride (CAS identifier 639-58-7) with Triphenyltin hydroxide (CAS identifier 76-87-9). **(c)** Weak MOA-cliff formed by Endosulfan sulfate (CAS identifier 1031-07-8) with Endosulfan I (CAS identifier 959-98-8).

## 4. Discussion and Conclusion

In this study, we explored and analyzed the activity landscape of chemicals in curated datasets of thyroid stimulating hormone receptor (TSHR) agonists (TSHR agonist dataset) and antagonists (TSHR antagonist dataset) compiled from the ToxCast library. By leveraging the established fingerprint-based approach and a substructure-based approach, we identified 79 activity cliffs in the TSHR agonist dataset and 69 activity cliffs in the TSHR antagonist dataset. Furthermore, we characterized the resultant activity cliffs based on the information on chemical structures. Additionally, we analyzed the differences in the mechanism of action (MOA) of the TSHR binding chemicals and identified 3 Strong MOA-cliffs and 19 Weak MOA-cliffs based on the difference in their annotated bioactivity outcomes. Notably, this is the first study to report the heterogeneity of the structure-activity landscape as well as the structure-mechanism relationships of the TSHR binding chemicals compiled from ToxCast chemical library.

The ToxCast library is among the largest datasets on environmental chemicals that have been assessed for their adverse biological effects (Dix et al., 2007; Richard et al., 2021). From our analysis, we also observed that the chemicals in the TSHR agonist and TSHR antagonist datasets have low chemical similarity among themselves, in particular, it has a median Tanimoto coefficient based similarity using ECFP4 fingerprints of ∼0.11. Further, we were unable to provide structure-based classification for the majority of the identified activity cliffs as the chemicals constituting these cliffs differ in their scaffolds as well as their scaffold topology (cyclic skeleton). Nevertheless, this identification of activity cliffs among the chemicals in ToxCast library will aid in development of robust toxicity predictors (Maggiora, 2006; Sheridan et al., 2020).

Activity cliffs arise when chemicals having similar structure show large differences in their activity values. We analyzed the activity cliffs in TSHR binding chemicals using two independent approaches, fingerprint-based structure-activity similarity (SAS) map and substructure-based matched molecular pair (MMP) approach. Interestingly, the activity cliffs identified via the MMP approach (MMP-cliffs) are a subset of those identified through the SAS map approach. This could be attributed to the highly restrictive fragment transformation conditions imposed in the generation of MMPs (Hu et al., 2012). Additionally, from chemicals common to both TSHR agonist and TSHR antagonist datasets, we identified 38 chemical pairs having high structural similarity, of which 9 chemical pairs (1 Strong MOA-cliff and 8 Weak MOA-cliffs) were also identified as activity cliffs through SAS map approach. Thus, these seemingly similar chemicals not only have differences in their binding affinity to TSHR, but also result in different downstream biological responses.

However, our workflow does not account for the stereoisomeric information of the chemical structures in identification of activity cliffs and MOA-cliffs. Moreover, we were unable to quantify the differences in binding affinities of chemicals forming MOA-cliffs as their affinity values are obtained from two different assays. We were also unable to explore molecular mechanisms behind the formation of activity cliffs as well as MOA-cliffs as there are no experimentally determined co-crystallized TSHR protein-ligand complexes available in the public domain.

Nonetheless, our efforts highlight the presence of activity cliffs and MOA-cliffs in a large chemical dataset such as ToxCast. In the future, one can use the newly developed chemical similarity methods such as extended similarity indices (n-ary comparison) (Miranda-Quintana et al., 2021a, 2021b) to deal with the computational complexity arising from pairwise comparison for large chemical datasets. In conclusion, this is the first investigation that combines SAS map and MMP approaches along with large-scale datasets from ToxCast chemical library to identify and characterize activity cliffs and MOA-cliffs among TSHR agonist and TSHR antagonist datasets. We believe that these insights will aid in development of better toxicity prediction models, and thereby, contribute towards characterization of the human exposome.

## Author Contributions

**Ajaya Kumar Sahoo:** Conceptualization, Data Compilation, Data Curation, Formal Analysis, Software, Visualization, Writing; **Shanmuga Priya Baskaran:** Conceptualization, Data Compilation, Data Curation, Formal Analysis, Software, Visualization, Writing; **Nikhil Chivukula:** Data Compilation, Data Curation, Formal Analysis, Writing; **Kishan Kumar:** Formal Analysis, Visualization; **Areejit Samal:** Conceptualization, Supervision, Formal Analysis, Writing.

## Acknowledgements

We thank Dhiraj Kumar for discussions. Areejit Samal acknowledges funding from the Department of Atomic Energy (DAE), Government of India and the Max Planck Society, Germany [Max Planck Partner Group in Mathematical Biology]. The funders have no role in the study design, data collection, data analysis, manuscript preparation, or decision to publish.

## Supplementary Information

### Supplementary Tables

**Supplementary Table S1:** Curated agonist dataset of 509 chemicals for thyroid stimulating hormone receptor (TSHR) from ToxCast chemical library. For each chemical, the table provides the CAS identifier or PubChem compound identifier (CID), chemical name, mechanism of action, modl_ga (logarithm of AC_50_ value in micromolar concentration) value, computed pAC_50_ value. Further the table provides chemical class information from ClassyFire, computed Bemis-Murcko scaffold using RDKit, documented endocrine disrupting chemical from DEDuCT, documented high production volume chemical from Organisation for Economic Co-operation and Development High Production Volume (OECD HPV) or United States High Production Volume (USHPV) databases, and documented OECD functional use(s) and Occupational health hazard report(s) from Chemical and Products Database (CPDat). Note that the modl_ga or pAC_50_ values are shown here up to 2 decimal places.

**Supplementary Table S2:** Curated antagonist dataset of 650 chemicals for TSHR from ToxCast chemical library. For each chemical, the table provides the CAS identifier or PubChem compound identifier (CID), chemical name, mechanism of action, modl_ga (logarithm of AC_50_ value in micromolar concentration) value, computed pAC_50_ value. Further the table provides chemical class information from ClassyFire, computed Bemis-Murcko scaffold using RDKit, documented endocrine disrupting chemical from DEDuCT, documented high production volume chemical from OECD HPV or USHPV databases, and documented OECD functional use(s) and Occupational health hazard report(s) from CPDat. Note that the modl_ga and pAC_50_ values are shown here up to 2 decimal places.

**Supplementary Table S3:** Activity cliff classification of 79 activity cliff pairs identified in the TSHR agonist dataset. For each chemical pair, the table provides the chemical identifiers, computed chemical similarity by using the Tanimoto coefficient between the ECFP4 fingerprints, activity cliff classification using the workflow (Figure 1d) and formation of MMP-cliff. Note that the Tanimoto similarity and the activity difference values are shown here up to 2 decimal places.

**Supplementary Table S4**: List of 9 activity cliff generators (ACGs) identified using the SAS map in the TSHR agonist dataset. For each ACG, the table provides the chemicals which form activity cliffs (separated by ’|’ symbol) and the number of activity cliff pairs.

**Supplementary Table S5:** List of 523 matched molecular pairs (MMPs) identified in the TSHR agonist dataset. For each MMP, the table provides the chemical identifiers, chemical SMILES (after performing canonicalization in RDKit and removing stereoisomer information) which was used then as input SMILES in mmpdb platform in order to generate MMPs, generated constant part and transformation between the chemical structures, and the activity difference between the chemicals (value shown here up to 2 decimal places). The chemical pairs forming MMPs and with activity difference of ≥ 2 (designated as MMP-cliff) are indicated in the last column.

**Supplementary Table S6:** Activity cliff classification of 69 activity cliff pairs identified in the TSHR antagonist dataset. For each chemical pair, the table provides the chemical identifiers, computed chemical similarity by using the Tanimoto coefficient between the ECFP4 fingerprints, activity cliff classification using the workflow (Figure 1d) and formation of MMP-cliff. Note that the Tanimoto similarity and the activity difference values are shown here up to 2 decimal places.

**Supplementary Table S7:** List of 4 activity cliff generators (ACGs) identified using the SAS map in the TSHR antagonist dataset. For each ACG, the table provides the chemicals which form activity cliffs (separated by ’|’ symbol) and the number of activity cliff pairs.

**Supplementary Table S8:** List of 590 matched molecular pairs (MMPs) identified in the TSHR antagonist dataset. For each MMP, the table provides the chemical identifiers, chemical SMILES (after performing canonicalization in RDKit and removing stereoisomer information) which was then used as input SMILES in mmpdb platform in order to generate MMPs, generated constant part and transformation between the chemical structures, and the activity difference between the chemicals (value shown here up to 2 decimal places). The chemical pairs forming MMPs and with activity difference of ≥ 2 (designated as MMP-cliff) are indicated in the last column.

**Supplementary Table S9:** List of 38 chemical pairs (which were common to both TSHR agonist and TSHR antagonist datasets, and have Tanimoto coefficient based similarity of > 0.35) and their mechanism of action (MOA) based classification. For each chemical pair, the table provides the chemical identifiers, computed chemical similarity by using the Tanimoto coefficient between the ECFP4 fingerprints (value shown here up to 2 decimal places), MOA annotations from both agonist and antagonist datasets and their MOA based classification as described in Figure 1e.

